# Briefer than brief - uncovering the true temporal resolution of the visual system

**DOI:** 10.64898/2026.08.20.745901

**Authors:** Joost de Jong, Claire Sergent, Mark Wexler

**Affiliations:** INCC, CNRS UMR 8002, Université de Paris Cité / CNRS, Paris, France

## Abstract

The temporal resolution of vision is seriously limited. However, the response to a very brief flash, called the impulse response, is already quite sluggish at the earliest stages of vision, potentially obscuring the true temporal resolution of the rest of the visual system. Faster monitors that produce briefer flashes are subject to diminishing returns because, by definition, they cannot elicit responses that are any briefer than the impulse response.

Here, taking inspiration from previous attempts, we develop a novel technique for presenting flashes that elicit ‘briefer-than-brief’ visual responses. Using a simple deconvolution technique, we reverse-engineer the visual response and estimate the form that the stimulus should take to elicit the response that a faster visual system would produce to a normal flash.

Using psychophysics on human observers, we demonstrate that these “briefer-than-brief” (BTB) flashes partially bypass the temporal limits presumably imposed by the early visual system using two paradigms: one that requires temporal segregation and one that requires temporal integration of sequential flashes. With BTB flashes, human observers successfully isolated two successive flashes at shorter intervals than with conventional flashes, improving temporal resolution by around 16%. We found that BTB stimuli not only improved temporal resolution, but also induced poorer performance on tasks requiring temporal integration, suggesting that the visual responses elicited by BTB flashes overlap less in time due to their briefer duration.

In sum, our findings suggest that, using reverse-engineered stimuli, we can alleviate a temporal bottleneck that probably originates from the earliest stages of vision. In doing so, we allow higher visual areas to operate at a higher temporal resolution than previously thought possible.

## Introduction

The temporal resolution of vision is seriously limited^1,2^. However, the response to a very brief flash, called the impulse response, is already quite sluggish at the earliest stages of vision^1,3,4^, potentially obscuring the true temporal resolution of the rest of the visual system^5^. Faster monitors that produce briefer flashes^6^ are subject to diminishing returns because, by definition, they cannot elicit responses that are any briefer than the impulse response.

Here, taking inspiration from previous attempts^7^, we develop a novel technique for presenting flashes that elicit ‘briefer-than-brief’ visual responses. Using a simple deconvolution technique, we reverse-engineer the visual response and estimate the form that the stimulus should take to elicit the response that a faster visual system would produce to a normal flash.

Using psychophysics on human observers, we demonstrate that these “briefer-than-brief” (BTB) flashes partially bypass the temporal limits presumably imposed by the early visual system using two paradigms: one that requires temporal segregation^8^ and one that requires temporal integration^9^ of sequential flashes. With BTB flashes, human observers successfully isolated two successive flashes at shorter intervals than with conventional flashes, improving temporal resolution by around 16%. We found that BTB stimuli not only improved temporal resolution, but also induced poorer performance on tasks requiring temporal integration, suggesting that the visual responses elicited by BTB flashes overlap less in time due to their briefer duration. In sum, our findings suggest that, using reverse-engineered stimuli, we can alleviate a temporal bottleneck that probably originates from the earliest stages of vision. In doing so, we allow higher visual areas to operate at a higher temporal resolution than previously thought possible.

### What limits visual temporal resolution?

Limits on visual temporal resolution have been well-documented^1,2^. For instance, flickering stimuli are perceived as continuously illuminated above 60 Hz^10^, and conversely, two successive flashes are seen as one when separated by less than 40 ms^11^. One major reason for this temporal limit is that the early visual system is sluggish^1,4,5,12^. Photoreceptors respond quickly to light, but they are slow in the sense that activity can linger for hundreds of milliseconds^1^. This sluggishness introduces significant temporal overlap between visual responses, thereby limiting visual temporal resolution.

Later stages in vision may also limit visual temporal resolution. For instance, some studies found that the frequency of cortical alpha oscillations partly determines whether two flashes are perceived as one integrated flash^11^. However, neurophysiological studies in primates suggest that most of the sluggishness of the visual system is due to photoreceptors^5,12^. This opens the possibility that later visual stages may operate at higher temporal resolution than previously assumed. This high temporal resolution in late visual stages may not be reflected in our subjective perception because of the sluggish bottleneck of the early visual system. Hence, visual perception could potentially operate at a finer temporal resolution if the early visual system were not so sluggish.

How can we get around the sluggishness of the early visual system? Producing ever briefer stimuli is not the answer, because even the briefest stimulus elicits a visual response that is spread out over time. Fortunately, we can use a reverse-engineering approach: the proper characterization of the visual impulse response permits us to create stimuli that elicit visual responses that are even briefer than the visual impulse response itself.

### Generating briefer-than-brief flashes

The temporal properties of the early visual system can be specified using an abstract visual impulse response function^13–16^. Theoretically, this is the response to an infinitesimally brief and infinitely strong pulse of light, a so-called Dirac delta. Practically, this amounts to the response to a flash. If we assume that the early visual system can be approximated linearly, the impulse response will tell us not just the visual response to a specified input, but also the reverse: what input is required to generate a specified visual response. Therefore, it is theoretically possible to generate visual responses that are much briefer than the visual impulse response^7,17^.

Taking inspiration from previous work^7^, we built the following approach for generating such Briefer than Brief, or BTB, stimuli (Figure 1). First, we use previous psychophysical findings^18,19^ to fix an impulse response function, with a temporal spread governed by the temporal parameter τ. This impulse response function determines the visual response to a brief Gaussian flash through convolution. Second, we compute what the visual response would look like if τ were smaller by some factor. This becomes our target response. Third, we compute the stimulus that produces this ‘briefer-than-brief’ target response given our *fixed* impulse response function through *de*convolution. This stimulus is a ‘briefer-than-brief’ stimulus, which is predicted to elicit a response that is briefer than the response to a conventional flash. Importantly, we can parametrically control how much the duration of the visual response is reduced by assuming that τ is reduced by an arbitrary ‘speed-up’ factor when computing the target response. In all experiments, we used a speed-up factor of 2, meaning that τ was reduced by 50% in calculating the target response. Importantly, this target response was briefer than the impulse response function itself (see Supplementary Figure 1).

**Figure 1.**
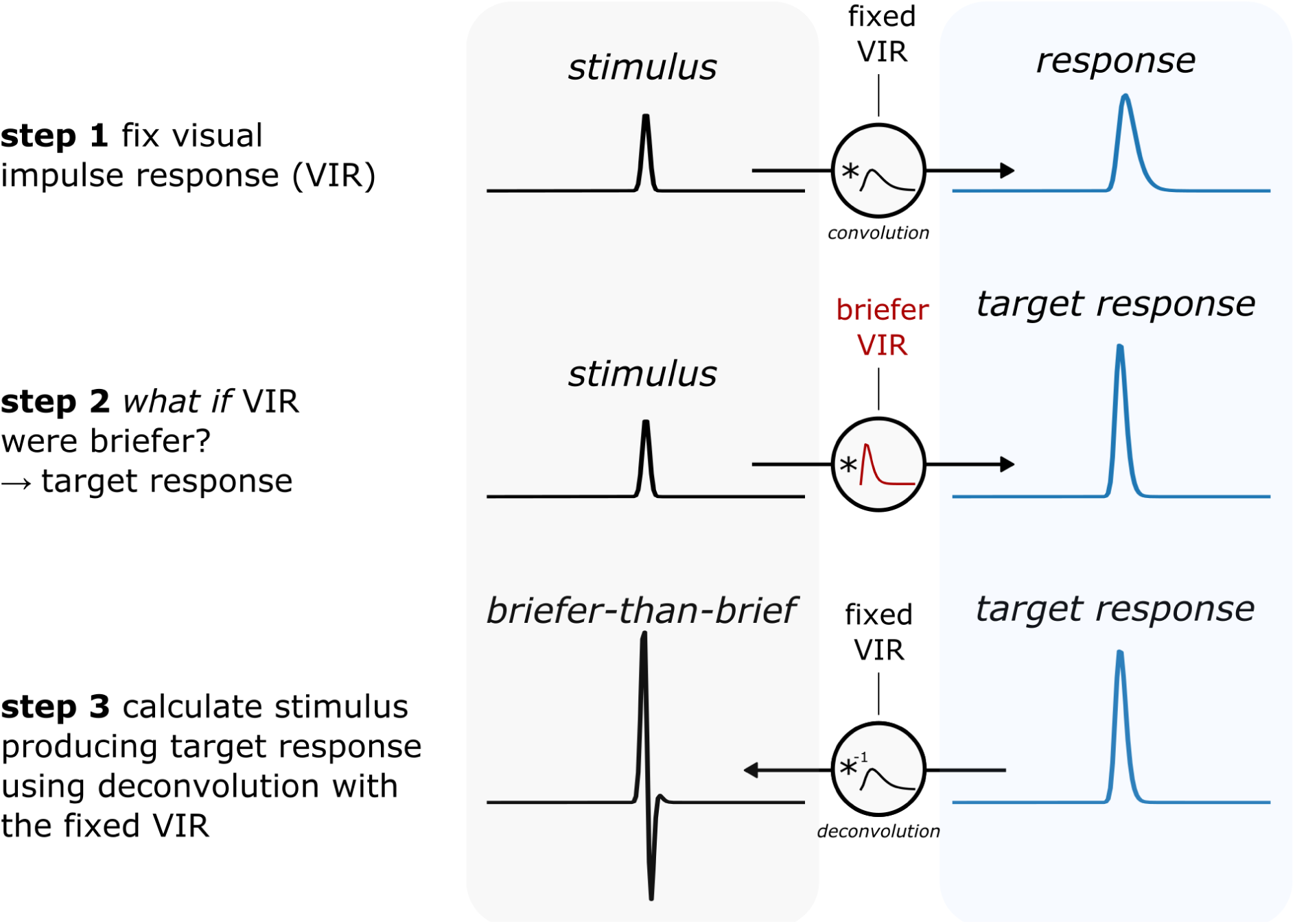
Approach for generating briefer-than-brief responses. Left panel: step-by-step explanation with pseudocode. Right panel, stimuli (in black) and visual responses (in blue) with the visual impulse response (VIR) determining their relation through (de)convolution. In step 1, we assume a VIR (see Methods) with τ governing its temporal spread. The visual response to an arbitrary stimulus (in this case a Gaussian flash) is determined through convolution. In step 2, we compute a ‘hypothetical’ VIR_b_, with τ_b_ reduced by a factor of s (i.e., τ_b_ = τ/s). This gives us what the visual response to an arbitrary stimulus would be if the VIR were briefer by some factor: the target response. In step 3, we use deconvolution to go from the target response to the stimulus that produces the briefer-than-briefer target response.

Our approach differs in some important ways from previous methods. In contrast to Tyler’s technique^7^, we estimate stimuli that *parametrically* reduce the temporal spread of the visual response, rather than minimizing it. In other words, our method allows us to speed up the visual response by a specific factor, instead of approaching an impulse-like visual response up to some order of approximation. Also, unlike a recently developed technique, our method is abstracted away from the physiological details of photoreceptors^17^, which means that it can be applied to any sensory system that can be approximated by a linear system.

Besides these methodological differences, previous studies did not actually test whether flashes designed to produce briefer-than-brief responses truly improve temporal resolution at the *perceptual* level. Discrete temporal windows at higher stages of vision may impose a strict limit on temporal resolution, regardless of how brief the neural responses are. Here, we experimentally test whether visual temporal resolution can be improved by presenting stimuli designed to generate briefer-than-brief responses.

### Briefer-than-brief flashes improve visual temporal resolution

We first measured visual temporal resolution using the two-flash paradigm (Figure 2). Participants (N=8) reported whether two successive flashes were perceived as a single flash or a flickering pattern of two flashes. The longer the interval between flashes, the more likely participants were to report a flicker. This stimulus-onset asynchrony (SOA) was varied according to a staircase procedure to maintain relatively constant performance. We used the resulting data to measure two-flash thresholds, the SOA at which participants reported seeing flickering 50% of the time, corrected for false alarms (reporting flicker when there was none). The lower the two-flash threshold, the better the visual temporal resolution. In the experiment, both conventional ‘Gaussian’ flashes and BTB flashes with a speedup of a factor of 2.0 were presented.

**Figure 2.**
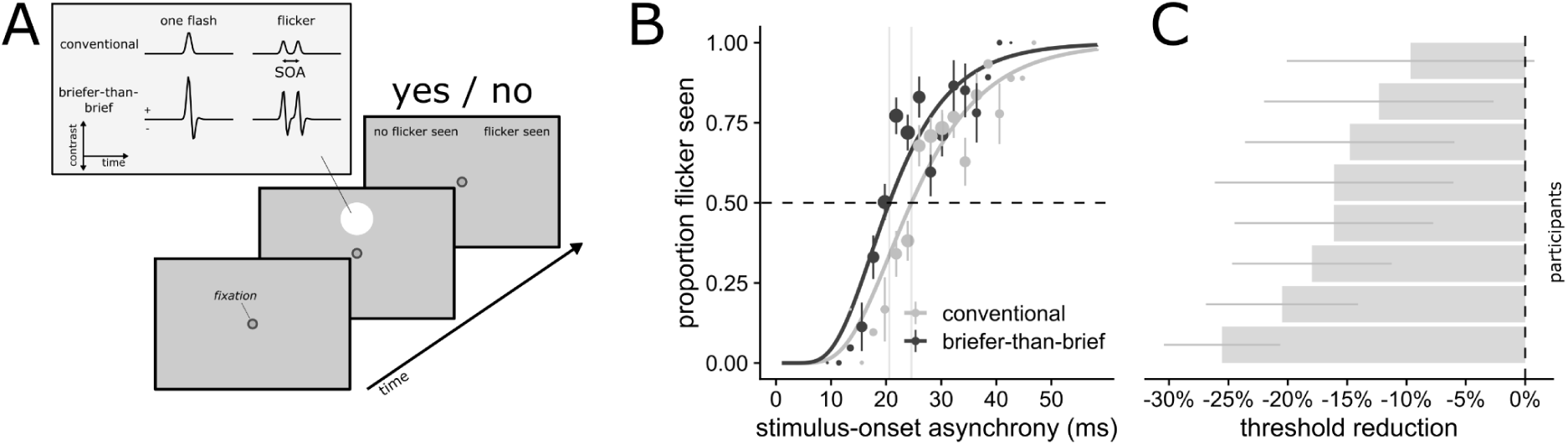
BTB stimuli improve visual temporal resolution in a yes-no task. (A) Stimuli were shown peripherally with luminance profiles computed with our method. Participants were presented with conventional or BTB stimuli that were either a single flash or a flicker, with a staircased stimulus-onset asynchrony (SOA). Participants responded whether they saw a flicker. (B) Proportion ‘flicker seen’ ± SE is plotted on the y-axis as a function of SOA on the x-axis. Individual points represent group-average proportions with associated standard errors, and their size represents the number of observations. Curved lines represent the fitted psychometric function. The horizontal dashed line represents threshold performance (50%), and vertical lines represent 2-flash thresholds. (C) Percent reduction in threshold for each participant with 95% confidence interval.

As shown in Figure 2B, all participants had lower two-flash thresholds for the BTB stimuli (M = 22.1 ms, SD = 4.7 ms) than for the conventional flashes (M = 26.6 ms, SD = 5.1 ms; *Z*-value = 8.59, dAIC = 77.88, dBIC = 72.73; Figure 2). On average, this amounted to a 16.6% reduction in threshold. To get a sense of how reliable this effect is at the individual level, we observed that six out of eight participants showed a statistically significant reduction in threshold (Holm’s adjusted p’s < 0.05), suggesting that BTB stimuli reliably improve visual temporal resolution at the individual level.

We ran some control experiments to assess the robustness of our technique. BTB stimuli, despite having the same integrated contrast as conventional stimuli, have more positive contrast and a higher peak contrast. Therefore, BTB stimuli may have higher apparent contrast and therefore induce response bias. Also, stimulus luminance may improve temporal resolution^20^, and could therefore partly explain improvements in visual temporal resolution. In one control experiment involving a single participant, we interleaved three contrast levels within a single session, thereby preventing contrast from serving as a cue for performing the task. We found that contrast improved temporal resolution, but our technique still worked for the two highest contrast levels (Supplementary Figure 2; 30% contrast: 3% increase, p = 0.888; 50% contrast: 19% reduction, p < 0.001; 80% contrast: 9% reduction, p = 0.002). Furthermore, the effectiveness of the BTB stimulus was also unlikely to be solely due to contrast, because BTB stimuli at 50% contrast had a lower threshold than conventional stimuli at 80% contrast (10% reduction, p < 0.001). Another potential issue is that decreases in contrast, similar to those in the BTB stimuli, may stimulate the OFF visual pathway, which has different dynamics than the ON visual pathway^21^. This may partly explain the efficacy of BTB stimuli, but also limit their applicability to arbitrary time-varying stimuli. However, in another control experiment, we found that our technique also worked for negative flashes (Supplementary Figure 3; 23% reduction, p < 0.001).

To investigate whether temporal resolution can be improved parametrically and whether these improvements can be pushed to the limit, two participants performed more extensive testing with multiple speed-up values. The target responses for the BTB stimuli corresponded to a temporal spread (τ) reduced by up to a factor of three (see Figure 3). They performed a 2-alternative forced-choice (2AFC) version of the experiment in which they reported which of two sequences contained a flicker, a paradigm that is less prone to bias. Two-flash thresholds decreased continuously with increased speedups, which was well approximated with an exponential function (R^2^ = 0.80; Figure 3B). The reduction in threshold asymptoted to around 26%, largely achieved at a speedup of 2.5 (Figure 3B). These findings suggest that the BTB technique can parametrically improve visual temporal resolution.

**Figure 3.**
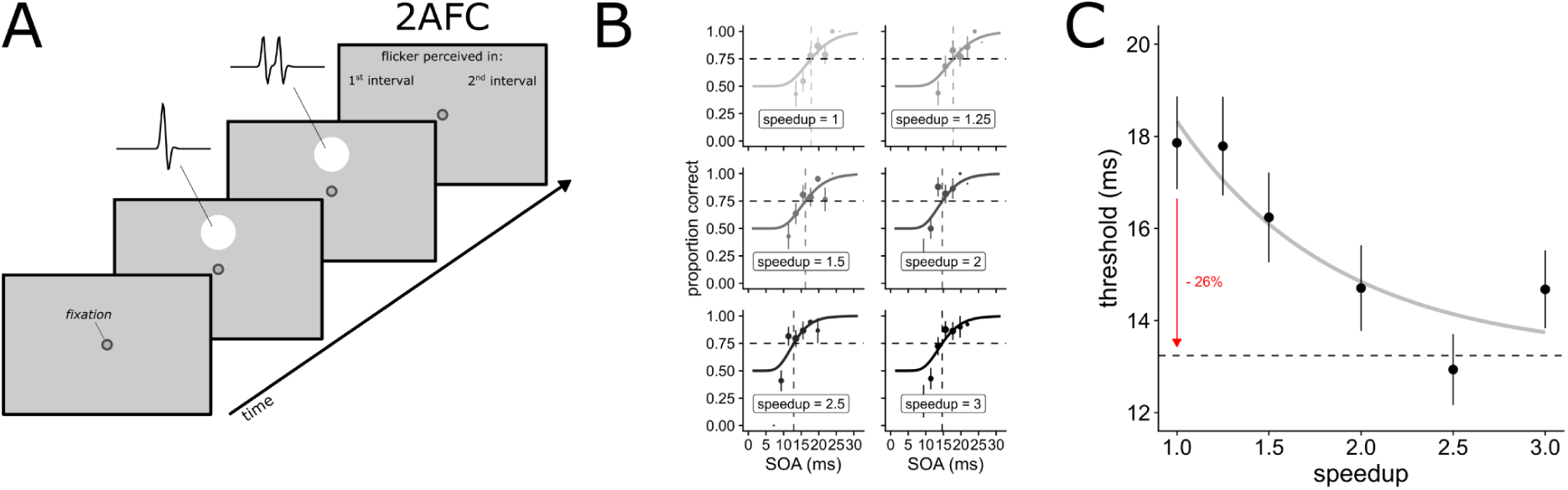
BTB stimuli parametrically improve visual temporal resolution in a 2AFC task. (A) On each trial, participants were presented with both single-flash and flicker stimuli in random order and reported which of the two contained the flicker. Both stimuli could be either conventional or BTB versions. (B) For each speedup, SOA is plotted on the x-axis, and the proportion of correct responses is plotted on the y-axis. The horizontal dashed line represents threshold performance (75%) and the vertical dashed line represents the threshold. (C) Threshold ± SE for different speedup factors. The continuous line represents a fit to an exponential function, and the dashed line represents its asymptote. The red arrow represents the percentage reduction from the threshold at no speed-up to the asymptote (26%).

Overall, our findings suggest that BTB stimuli improve visual temporal resolution substantially, reliably, and parametrically. It is not clear, however, whether this improvement is specific to the task measuring temporal resolution, as intended, or whether BTB stimuli unintentionally boost task performance in general. We reasoned that if BTB stimuli generate ultra-brief responses, they must have a flip side: they should degrade performance on a task that requires integrating signals over time.

### Briefer-than-brief flashes impair visual temporal integration

Vision integrates information over time to form a coherent impression of the outside world^22^. A common assumption is that temporal integration is driven by the temporal overlap or correlation between visual responses^23,24^. Therefore, if BTB stimuli produce briefer visual responses with less temporal overlap, they should undermine temporal integration.

In order to test this prediction, we assessed temporal integration^9^ using the ‘missing element’ paradigm. Two consecutive spatial patterns were flashed, which, when integrated, together formed a circle with a single missing element that participants needed to locate (Figure 4A). Thus, the performance of locating the missing element is a measure of temporal integration. As expected, we found that, as the stimulus-onset asynchrony (SOA) increased, participants (N=8) were less able to locate the missing element (z = 15.08, p < 0.001; see Figure 4B). Crucially, this drop in performance happened at shorter SOA for BTB stimuli compared to conventional stimuli (z = −4.51, p < 0.001), demonstrating that they impair temporal integration. At the individual level, we found a statistically significant effect for only half of the participants; however, the aggregate reduction in threshold in the missing element task (18%; p = 0.001) was remarkably similar to that observed in the two-flash paradigm (16% reduction).

**Figure 4.**
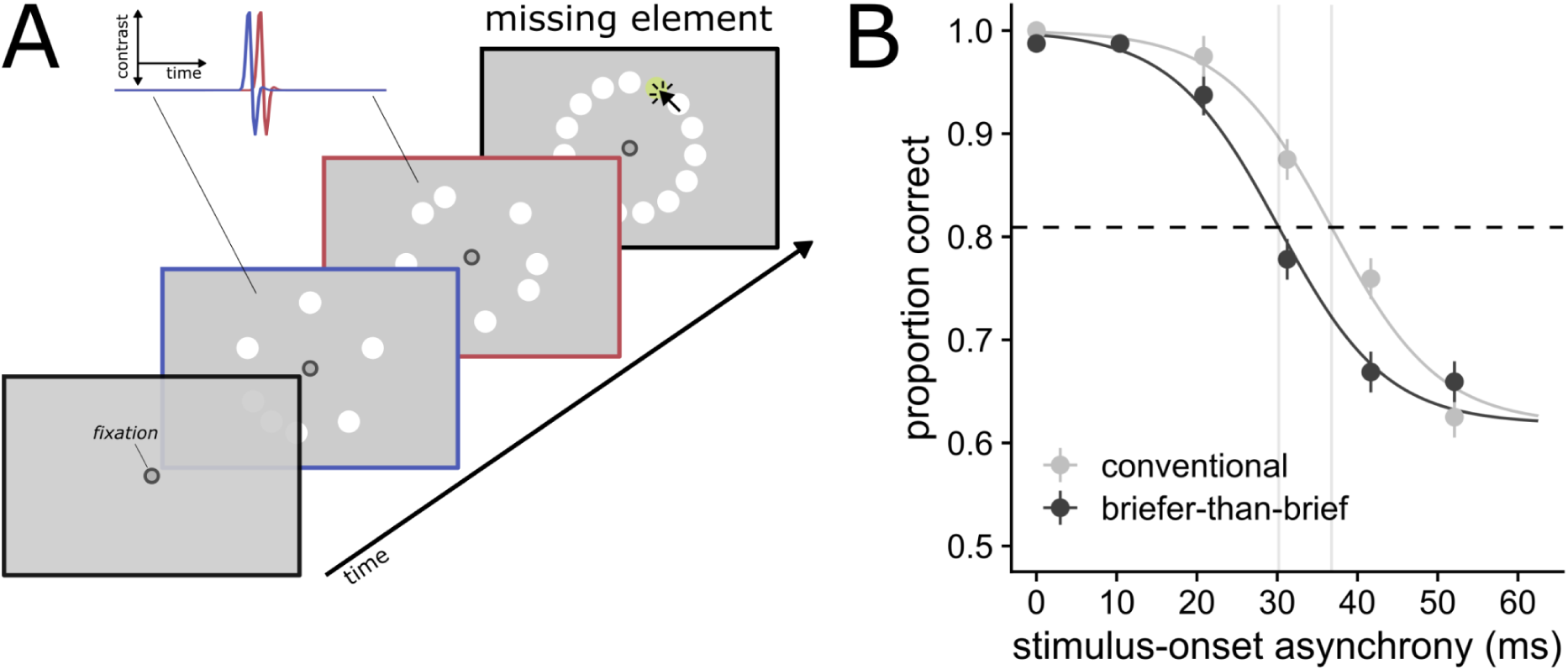
BTB stimuli impair temporal integration. (A) Two circular arrays of dots were sequentially flashed, either conventionally or BTB, with a stimulus-onset-asynchrony (SOA) ranging from 0 to 52 ms. The union of these arrays formed a circle of dots with one missing element (in green), which participants needed to locate using a mouse. (B) Proportion correct ± SE is plotted on the y-axis as a function of SOA. Smooth curves are psychometric curves with a logistic link ranging from 1.00 to a fitted asymptote of 0.62. The horizontal dotted line represents threshold performance (halfway point between the two asymptotes) and vertical lines represent the thresholds for conventional and BTB stimuli (crossing point between the psychometric curve and threshold performance).

Overall, these findings suggest that BTB stimuli produce visual responses that are substantially briefer than responses to their conventional counterparts, and therefore not only improve temporal resolution but also impair temporal integration.

## Discussion

Vision has clear limits on temporal resolution. However, because these limits can be largely attributed to the temporal spread of the impulse response at early visual stages, it is not clear whether the rest of the visual system is similarly limited.

Here, we developed a technique for displaying flashes in a way that produces early visual responses briefer than those to a flash. Our findings show that these ‘briefer-than-brief’ flashes substantially improve visual temporal resolution, suggesting that the rest of the visual system may operate at a much higher temporal resolution than previously thought.

Leveraging the parametric nature of our method, we were also able to improve visual temporal resolution parametrically. In other words, BTB stimuli can, up to a limit, parametrically control the temporal resolution of vision. BTB flashes also improve temporal resolution across a range of contrasts and for flashes formed by luminance decrements, which suggests that they can be applied to arbitrary time-varying stimuli. We also found the flip side to this improvement in temporal resolution: BTB flashes degraded temporal integration. Overall, these findings suggest that BTB stimuli elicit visual responses substantially shorter than those to flashes, and therefore not only improve temporal resolution but also impair temporal integration.

Our novel BTB technique can be applied and extended in several ways. First, BTB stimuli can be applied to other temporal percepts, such as temporal order, simultaneity, or duration. Studies like this can uncover whether the sluggishness of early sensory stages generally limits temporal perception, or whether later visual stages impose a more discrete limit^25^. Furthermore, the reduced temporal spread of neural responses induced by BTB stimuli may allow neuroscientists to disentangle processes that normally overlap too much in time. Moreover, given its linear assumptions, visual responses can be sped up for arbitrary time-varying stimuli, such as naturalistic videos. Finally, to the extent that the early stages of a sensory system, such as the auditory system^26^, can be approximated linearly, our method may also generate BTB sounds that could benefit speech comprehension in sped-up audio.

There are some potential limitations to our technique. The reliability of BTB stimuli is only as good as our model of the impulse response function. The early visual system is not strictly linear, and visual responses may speed up at higher contrasts^27^. However, within a range of contrasts, vision operates nearly linearly^28^. Even though our method seems to work across a range of contrasts (Supplementary Figure 2), it is difficult to estimate the extent to which non-linearities affect it.

A more tractable violation of our assumptions is parametric: What if the real value for the temporal spread of the impulse response function is smaller or larger than assumed? To quantify how efficiently BTB stimuli generate sped-up visual responses, we generated BTB stimuli at various speedups at a fixed value of τ, then computed the visual response through convolution with a VIR that had a higher or lower τ, and finally recovered τ given the original (non-BTB) stimulus and the visual response. Intuitively, this procedure exactly recovers the speedup factor when τ is accurately specified. Using this ‘parameter recovery’ procedure, we found that if the real τ is over- or underestimated by a factor of 1.5, speed-ups still have more than 50% efficiency (Supplementary Figure 4). This suggests that our technique is relatively robust to parameter misspecification, but also that even larger improvements in visual temporal resolution are possible with more accurate models of the impulse response function, calibrated to differences between stimuli and individuals.

In sum, using a simple deconvolution technique, we were able to generate ‘briefer-than-brief’ stimuli that elicit visual responses briefer than the response to a conventional flash. We found that BTB stimuli parametrically improved visual temporal resolution across a range of conditions, but also impaired temporal integration. Our BTB method demonstrates that higher visual areas may operate at a faster timescale than previously thought, and it suggests a straightforward way to study and drive perception at unprecedented temporal resolution.

## Acknowledgments

We wish to thank Christopher Tyler and Patrick Cavanagh for useful discussions. This work was supported by Agence Nationale de la Recherche grant ANR-22-CE28-0025.

## Code and data availability

Python code for generating BTB stimuli and parameter recovery is available on GitHub: https://github.com/dejongejoost/briefer-than-brief. Behavioural data from experiments are available on Zenodo: https://doi.org/10.5281/zenodo.21903986

## Methods

### Generating briefer-than-brief stimuli

Here, we outline our technique for generating briefer-than-brief (BTB) stimuli in more detail. The technique consists of three steps:

1. Define the visual impulse response function (VIR) and compute the response (R) to a Gaussian flash stimulus (S) through convolution (⊗).
2. Compute the ‘target response’ (R_target_). This is the response to a Gaussian flash *if* the impulse response’s duration (governed by τ) were briefer by some speedup factor (*s*).
3. Compute the stimulus (S_btb_) that produces the BTB target response through deconvolution (⊗^−1^) with the original impulse response function (VIR).

#### Working Python code for generating BTB stimuli

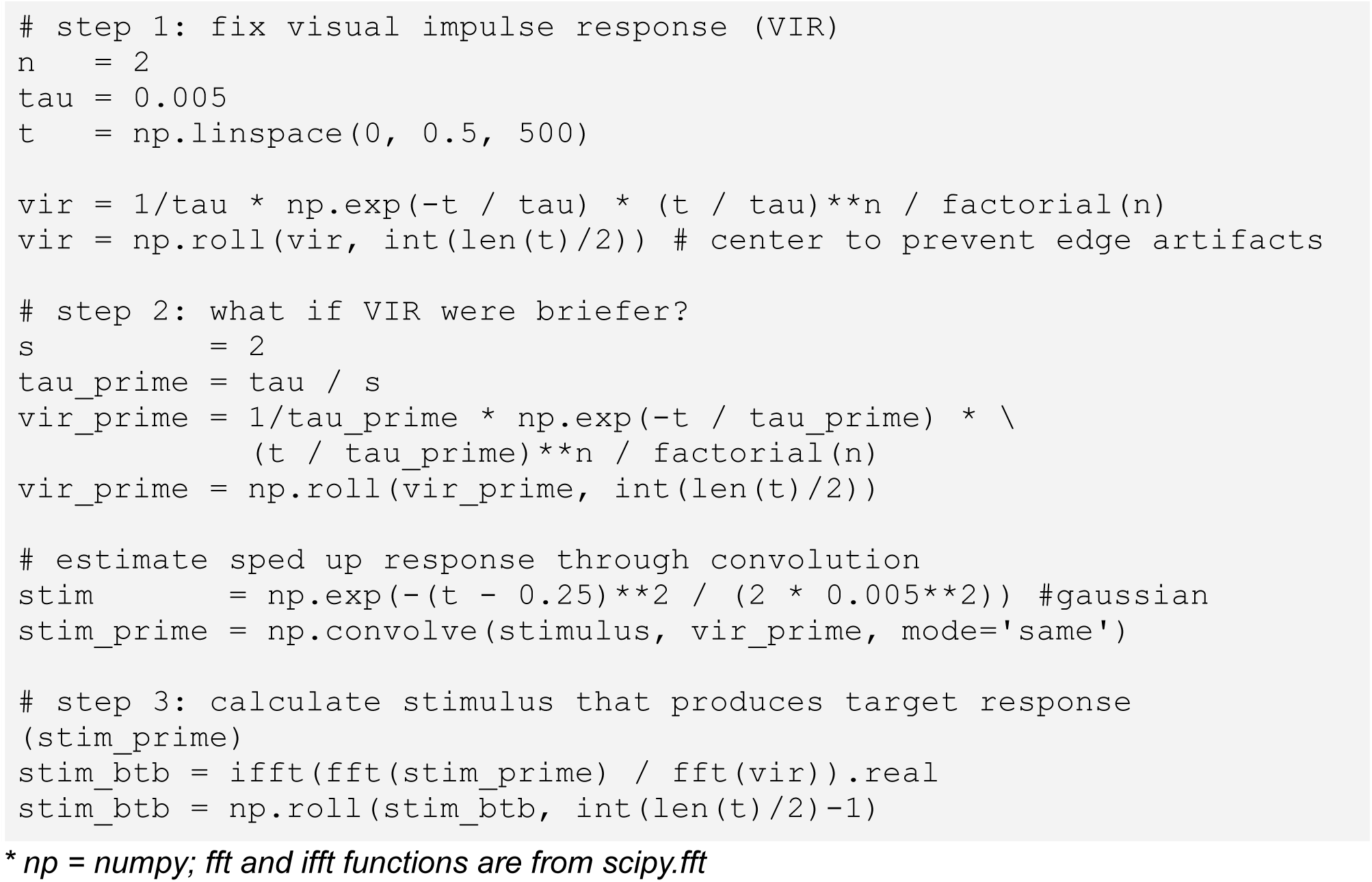

#### Step 1: Define the visual impulse response function

First, we need to specify how the early visual system responds to a very brief flash. This visual impulse response function (VIR) can take different forms, but we have opted for a relatively simple monophasic version of the VIR^18^, which is derived from psychophysical data^19^:

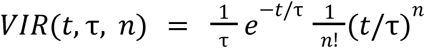

where *n* equals 2 and τ equals 5 ms. The parameter that controls the temporal spread, τ, was determined by initializing it so that the negative contrast phase of the BTB stimulus could be clearly seen by the first participant (first author of this paper). Then, τ was gradually reduced until the negative phase could no longer be seen (i.e., the BTB stimulus was perceived as a single, positive contrast flash), and this value was used for all experiments and all participants.

We then compute the response to a ‘Gaussian’ flash stimulus (S), which is an impulse convolved with a Gaussian kernel (σ = 3 ms). This step is necessary because it filters out high frequencies in the target response (Step 2). If these high frequencies are still present in the target response, the BTB stimulus needs to have prohibitively high contrast to produce a visible response.

#### Step 2: Compute BTB target response

Second, we compute the response that our stimulus should generate, the target response (R_target_). In principle, the target response can be any time-varying signal, such as an impulse or a step function. However, here we take a parametric approach, which ensures that any arbitrary time-varying stimulus could be made to produce faster responses than normal. R_target_ is the visual response to the Gaussian flash that *would* be produced if τ were reduced by a factor of *s* (τ/*s*). Hence, the target response has the same form as the VIR in every respect but its temporal spread, making it briefer than the normal visual response to the Gaussian flash.

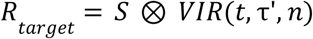

where τ′ equals τ/*s*.

#### Step 3: Compute stimulus producing BTB target response

Finally, we use the assumed VIR (with the original value of τ) to estimate which stimulus would produce R_target_ through deconvolution:

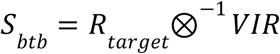

But it is most straightforward to implement this in the frequency domain:

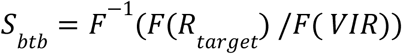

where *F*(·) denotes the Fourier transform and F^−1^(·) denotes the inverse Fourier transform. The resulting *S_btb_* produces the target response when it is convolved with our assumed VIR (with the original value of τ).

It is important to note that the BTB stimulus is generated not just by boosting the amplitude of high-frequency components. Our technique also shifts the *phases* of the frequency components, which compensates for the phase shifts induced by the VIR, in order to produce a target response (Supplementary Figure 5). Therefore, improvements in visual temporal resolution by BTB stimuli are not only due to higher frequency content in the stimulus, but also precise phase adjustments. Another important point is that we are not actually speeding up the VIR or the underlying dynamical mechanisms that produce the VIR. Instead, our technique leverages the VIR in order to generate visual responses that *would* have been produced with a ‘hypothetical’, briefer VIR (i.e., with a smaller τ).

### Two-flash paradigm

#### Participants

Participants (N=8; age 26 - 58y; 3 female) were the authors and colleagues of the authors who volunteered to participate in the experiment. Informed consent was obtained before participation. The study was approved by the CPP Ouest V ethics committee (dossier 2021-A00815-36).

#### Apparatus and Stimuli

Stimuli were presented on a 2560 x 1440 OLED monitor (SDM27Q10S*10, SONY) running at 480 Hz and driven by an NVIDIA RTX A1000 GPU. This OLED monitor, similarly to other recent high-speed OLED monitors, has extremely fast rise and fall times, allowing the presentation of fine temporal detail. The luminance of the monitor was manually calibrated using a chroma meter (Minolta CS-100). The experiment was programmed in OpenSesame 4.1.0 ^29^.

All stimuli were presented against a grey background at around 35.5 cd/m^2^. Participants were instructed to fixate on a central fixation dot that was present throughout the experiment. Both conventional and BTB flashes were circles with a size of 1.04° and presented 2.5° above the fixation dot. Flashes were presented by changing the luminance of the circle on each refresh (i.e., each 1000ms / 480Hz = ~2.1ms) according to the luminance profiles computed by our technique, where values of zero corresponded to the background luminance of 35.5 cd/m^2^. In all experiments, unless stated otherwise, stimuli were presented at 50% contrast. Participants were seated in a sound-attenuated dark room in a comfortable chair, with their head on a chin rest approximately 55 cm from the screen.

#### Yes-No psychophysical procedure

In all two-flash experiments, we used the yes-no psychophysical procedure, unless stated otherwise. On each trial, participants were presented with one out of four possible stimuli (equal probability): (1) a single conventional flash, (2) a single BTB flash, (3) two conventional flashes with a stimulus-onset asynchrony (SOA), or (4) two BTB flashes with an SOA. Trials started with a blank period of 500 ms, after which the stimulus was presented (see Figure 2).

Participants responded with keyboard buttons whether they perceived a flickering pattern of two flashes (right key: ‘yes’) or a single flash (left key: ‘no’), without a time limit. The SOA between the double flashes was varied according to the SIAM staircase procedure^30^, an unbiased method for estimating perceptual thresholds using yes-no tasks. Separate staircases were run for conventional and BTB stimuli, both starting at an SOA of 41.7 ms. When participants made a correct judgment, the SOA was decreased by one step (2.1 ms), making the task more difficult; when they made an incorrect judgment, the SOA was increased by three steps (6.3 ms), making the task easier. This weighted up-down rule^31^ converges to an accuracy level of 75%, which it also did in our experiment. Participants performed 320 trials with a self-paced break every 40 trials.

In the experiment where contrast was randomly varied (N=1), stimulus contrast was 30%, 50%, or 80%, and separate staircases were run for each unique contrast and stimulus type (conventional vs. BTB). In the negative contrast experiment (N=1), the contrast of the Gaussian stimulus was flipped, such that the flash consisted of a decrease in luminance. The BTB stimulus was computed in exactly the same way as the other BTB stimuli, as described in the section ‘*Generating BTB stimuli’*, which amounted to a contrast-flipped version of the BTB stimulus.

#### Two-alternative forced-choice (2AFC) procedure

To collect converging evidence that our BTB stimuli improve temporal resolution, and test whether this can be done parametrically, we employed the 2AFC paradigm. The 2AFC paradigm is less bias-prone than the yes-no paradigm, since participants do not report whether a flicker was presented (which depends on the participant’s criterion for reporting flicker) but which of two sequentially presented stimuli contains a flicker.

The 2AFC experiment (N=2) was identical to the yes-no experiments in terms of the apparatus and stimuli. On each trial, participants were presented with two consecutive stimuli, with one of them containing a single flash and the other a flicker with a staircased SOA. Participants responded with mouse clicks whether the flicker was presented first (left button) or second (right button). The SOA was staircased with the same procedure as the yes-no task: SOA was decreased 2.1 ms after a correct response and increased 6.3 ms after an incorrect response, which converges to an accuracy level of 75%. In the 2AFC experiments, instead of using a single speedup factor of 2, we ran separate staircases for speedup factors 1.0 (which is the original Gaussian stimulus) to 3.0 (which amounts to a reduction in τ of 66%).

#### Analysis

We used R (version 4.4.2) for the behavioural analyses. All analyses were performed both at the group level to assess the overall efficacy of the BTB stimuli and at the individual level to assess their reliability.

We fitted a Generalized Linear Model (GLM) with the binary yes/no response as the dependent variable, with only trials in which a flicker was presented. We used a probit link function that ranged from 0.00 to 1.00 for the yes-no task and from 0.5 to 0.999, using the probit.2asym function from *psyphy* (version: 0.3), for the 2AFC task. To correct for differences in false-alarm rate between conventional and BTB stimuli, we introduced an offset term in the model, which is the difference in z(FA) between the conditions^32^. We computed z(FA) using the *loglinear* approach^33^ to account for false-alarm rates near zero (FA = (N_false-alarms_ + 0.5) / (N_no flicker_ + 1)).

To statistically assess whether BTB stimuli reduce 2-flash thresholds, we used model comparisons. In the null model, the only predictor was log(SOA). Stimulus type (conventional vs. BTB) was added as a categorical predictor, with conventional as the reference level. We report the z-value for this predictor and the difference in AIC (dAIC) and BIC (dBIC) between the null model and the model containing stimulus type.

We quantified the magnitude of the effect by computing the threshold, which is the SOA at which the probability of responding ‘flicker seen’ is 0.5 for the yes-no paradigm and 0.75 for the 2AFC paradigm, for both the conventional (Δ*SOA_conv_*) and BTB stimulus (Δ*SOA_btb_*):

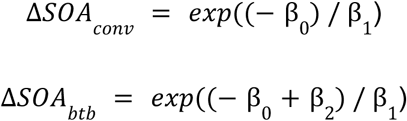

where β_0_ is the intercept, β_1_ is the coefficient for *log*(*SOA*), and β_2_ corresponds to the coefficient for the BTB stimulus type. Their associated 95% confidence intervals were computed using the Delta Method for both conventional and BTB stimuli. The reduction in threshold (in percentage; (Δ*SOA_conv_* − Δ*SOA_btb_*) / Δ*SOA_conv_*) and its 95% confidence interval were also calculated using the Delta method. Pairwise comparisons were performed using the *emmeans* package with default settings (version: 1.11.0). The exponential decrease in threshold with increasing speedup was done using nls.multstart (version: 1.3.0).

### Temporal integration paradigm

#### Participants

Participants (N=8; age 24 - 58 y; 3 female) were the authors and colleagues of the authors who volunteered to participate in the experiment. Informed consent was obtained before participation. The study was conducted in accordance with […] and approved by the […].

#### Apparatus, Stimuli, and Procedure

We used the same apparatus as in the previous experiments, but in order to reliably draw 15 circles in time for each screen flip (2.3 ms), this experiment was programmed using custom Python code that interfaced with OpenGL.

Stimuli were presented against a grey background at around 35.5 cd/m^2^. Participants were instructed to fixate on a central fixation dot that was present throughout the experiment. Two arrays of seven dots each (size = 2.40°; eccentricity = 4.16°) were presented simultaneously (SOA = 0 ms) or sequentially (SOA > 0 ms), together forming a circular array of dots with a single missing element. Then, a response screen with the complete circular array was presented, and participants clicked on the element that was missing. Participants were instructed to try to determine the exact location of the missing element, but that being off by one circle would be considered good performance. The missing element always lit up green after the response, and additionally, after an incorrect response, the clicked circle turned red. The arrays with eight circles were presented by changing the luminance of the circles on each refresh (i.e., each 1000 ms / 480 Hz = ~2.1 ms) according to the luminance profiles computed by our technique, where values of zero corresponded to the background luminance of 35.5 cd/m^2^. Participants were seated in a sound-attenuated dark room in a comfortable chair, with their head on a chin rest approximately 55 cm from the screen. Participants performed 480 trials with a self-paced break every 40 trials.

#### Analysis

We used R for the behavioural analyses. All analyses were performed both at the group level to assess the overall efficacy of the BTB stimuli and at the individual level to assess their reliability.

We fitted a Generalized Linear Model (GLM) with accuracy (whether the response was within one circle distance of the missing element) as the dependent variable, and SOA and stimulus type (conventional vs. BTB) as independent variables. We used a logit link function (using the logit.2asym function from *psyphy* (version: 0.3), with an upper asymptote of 1.00 and an estimated lower asymptote (using the ‘optim’ function in R, by minimizing the GLM’s AIC). Threshold performance was defined as the halfway point between the upper and lower asymptotes, and we used the ‘optim’ function to find the threshold, i.e., the SOA at which the fitted psychometric curve reached threshold performance. The reduction in threshold in percentage equals (Δ*SOA_conv_* − Δ*SOA_btb_*) / Δ*SOA_conv_*. To assess whether this reduction in threshold was statistically significant, we performed a permutation test by randomly shuffling condition labels (conventional vs. BTB) 1,000 times.

## Supplementary figures

**Supplementary Figure 1.**
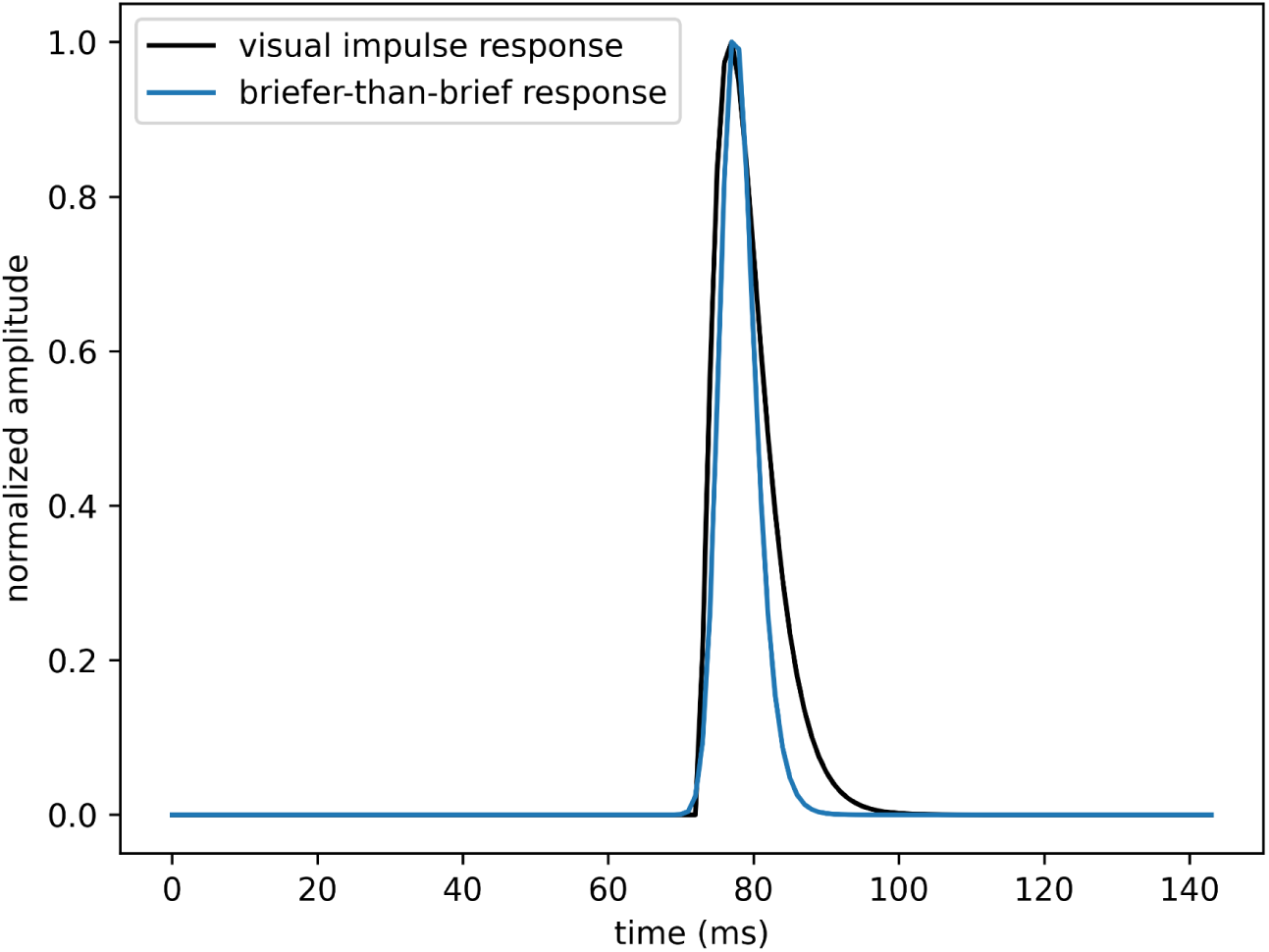
BTB response (speedup = 2.0) plotted alongside the visual impulse response from Figure 1. The BTB visual response (in blue) is briefer than the visual impulse response (i.e., the response to the briefest possible input; in black).

**Supplementary Figure 2.**
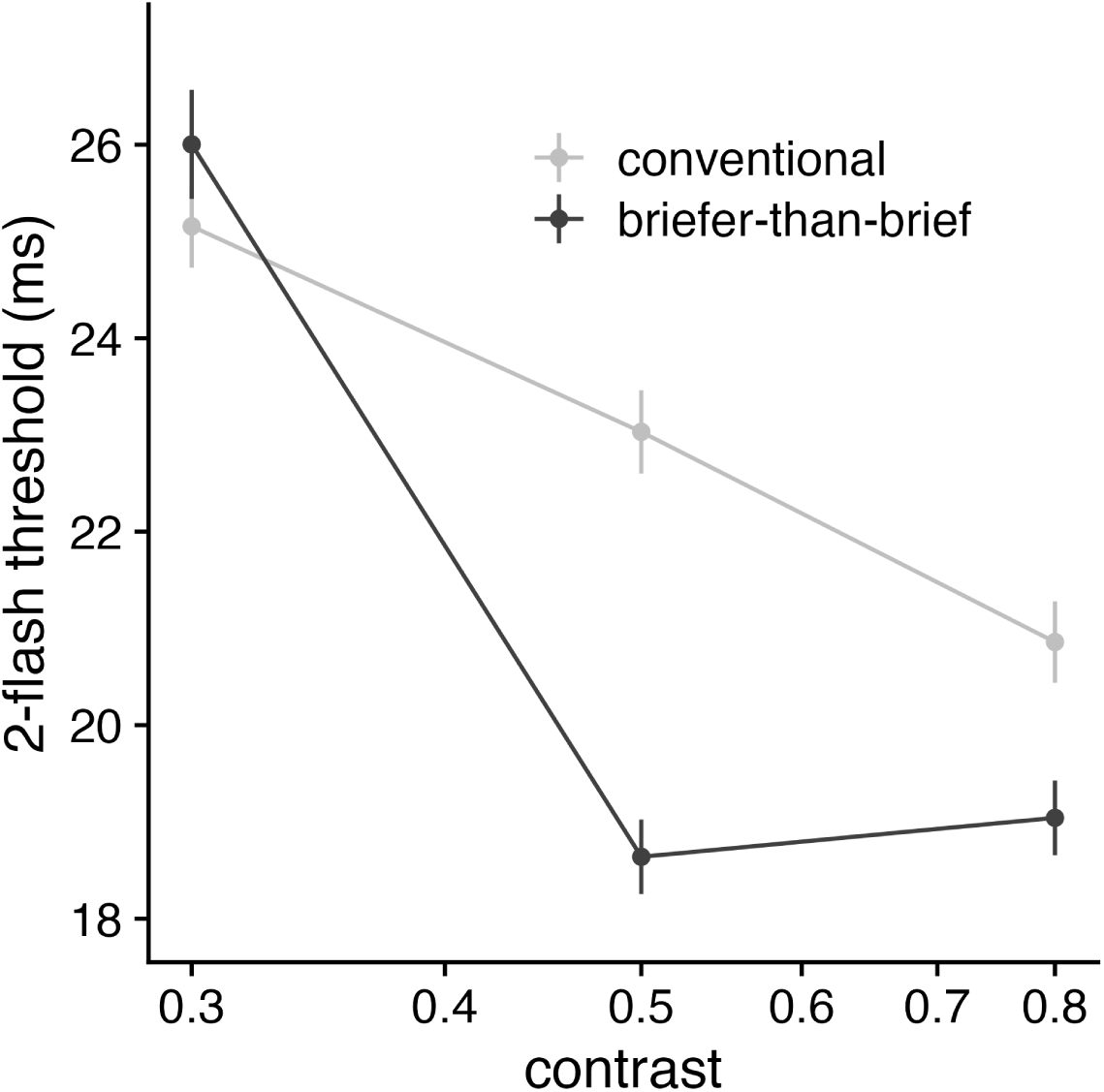
2-flash thresholds (±SE) for conventional and BTB flashes across three levels of stimulus contrast.

**Supplementary Figure 3.**
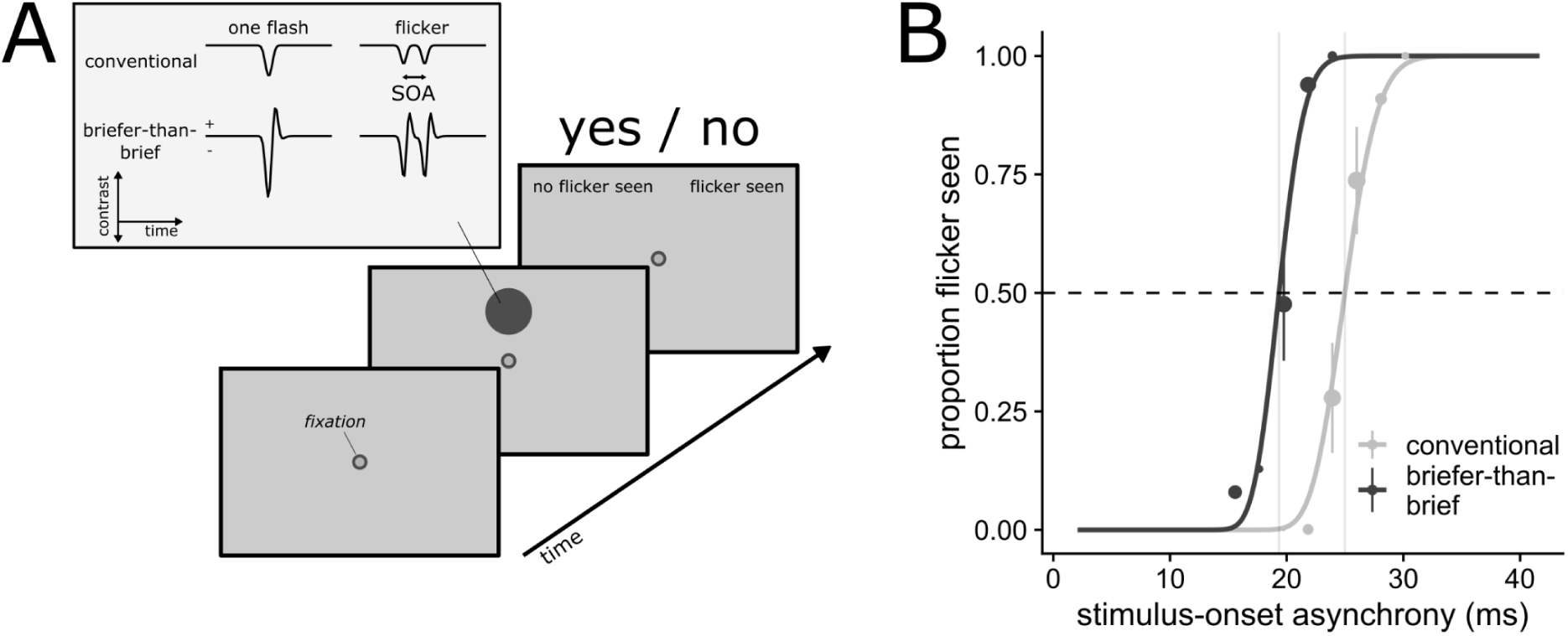
BTB with ‘negative flashes’, which consist of luminance decrements. A: experimental design uses the staircased yes/no paradigm, but with contrast flipped stimuli. B: proportion of flicker seen as a function of SOA for conventional and BTB stimuli.

**Supplementary Figure 4.**
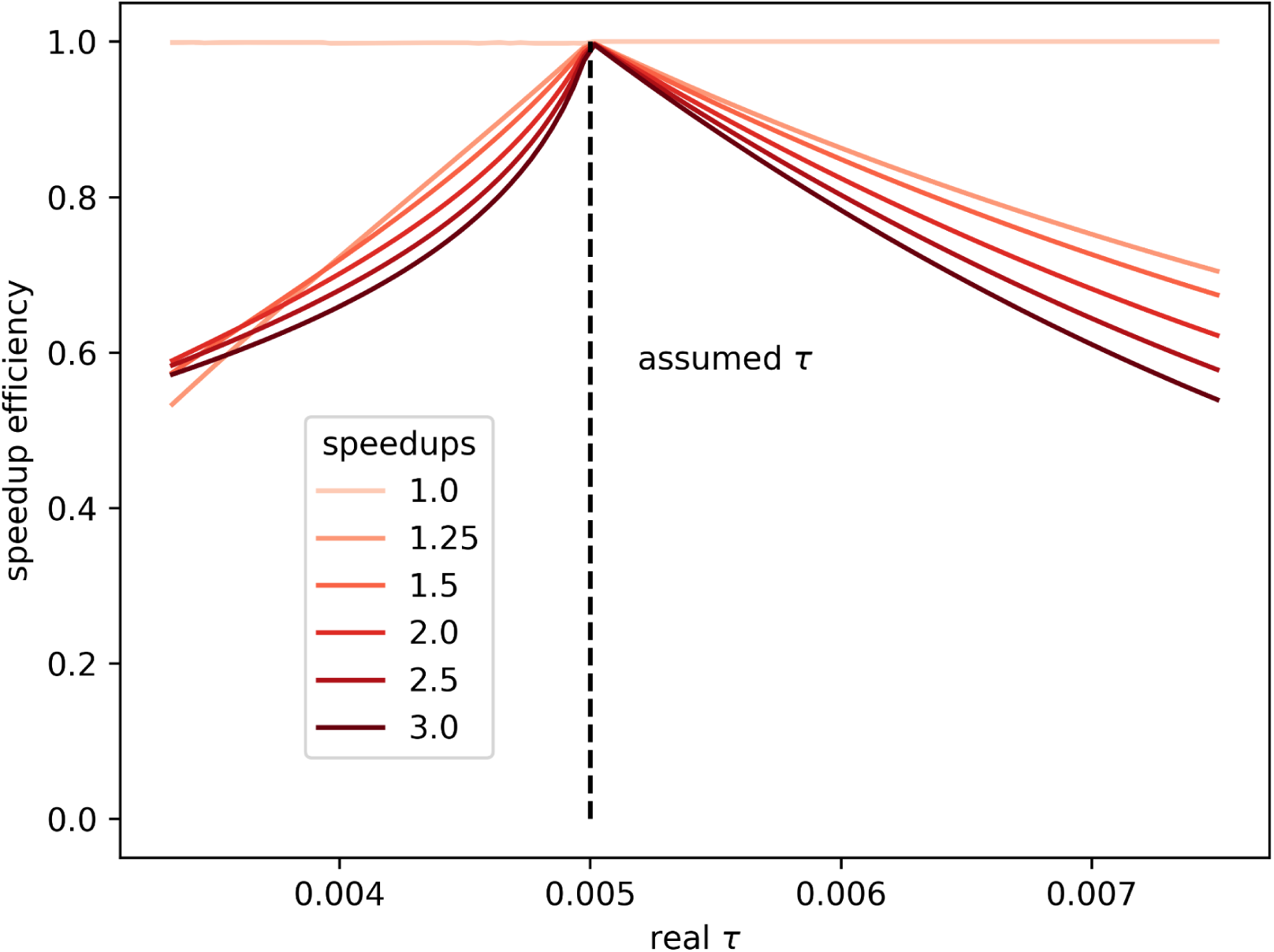
Efficiency estimates for BTB stimuli using parameter recovery. The BTB stimulus was computed with an assumed τ = 5 ms at various speedups. These stimuli were convolved with a ‘real’ VIR that had smaller or larger τ, resulting in a ‘real’ visual response. The effective speedup of this visual response was estimated by finding the τ that best fitted the visual response to the Gaussian stimulus. When the assumed τ matches the real τ, the estimated reduction in τ exactly matches the speed-up (efficiency = 1.0). When the intended speedup is 2.0, but the achieved speed-up is 1.5, efficiency equals 0.5.

**Supplementary Figure 5.**
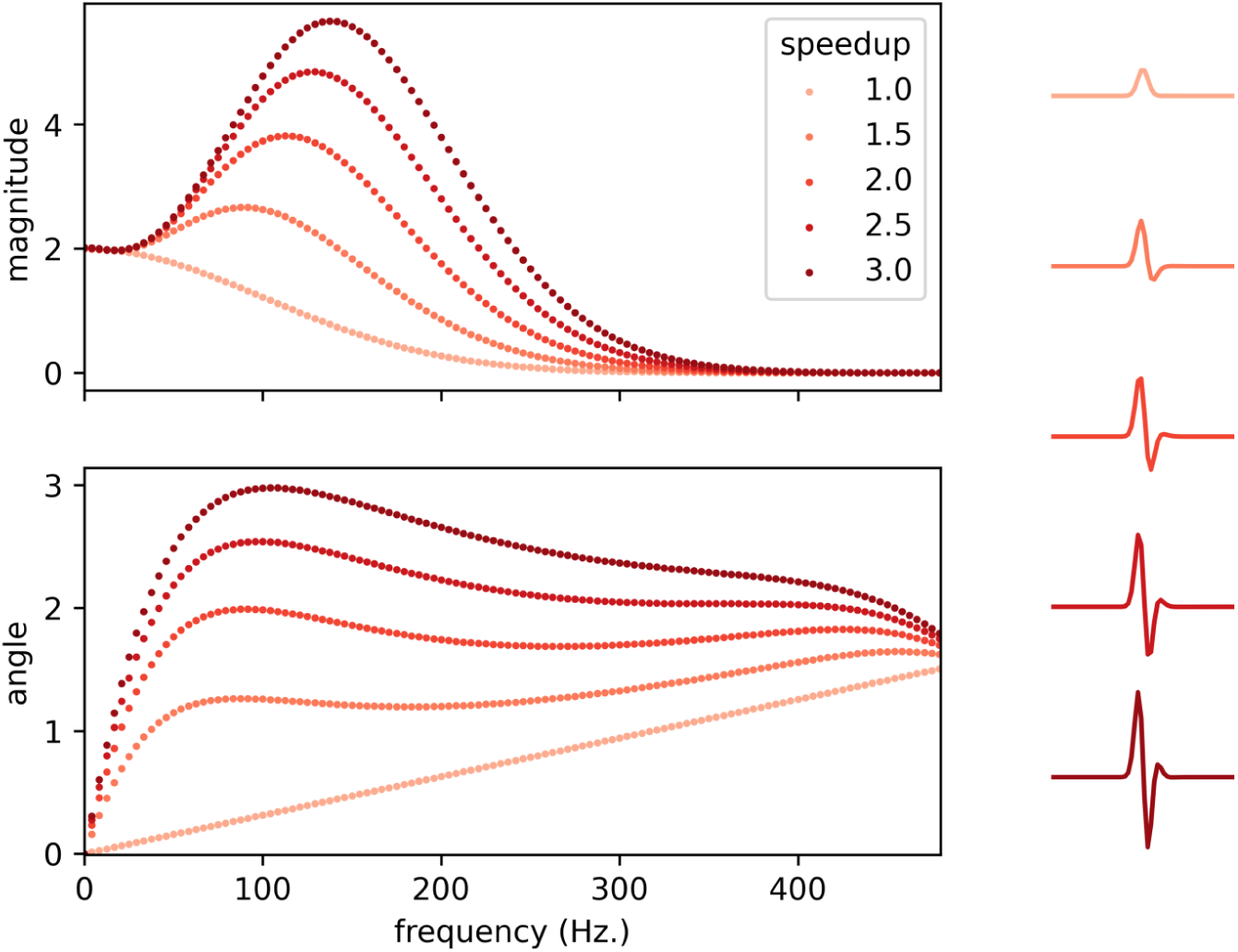
Magnitude and phase modulation of BTB stimuli. The magnitude (top panel) and angle (bottom panel) of each sinusoidal frequency component across different speedups. Time-domain BTB stimuli are plotted on the right.

